## Supplemental figures for "A natural variant of the sole pyruvate kinase of fission yeast lowers glycolytic flux triggering increased respiration and oxidative-stress resistance but decreased growth"

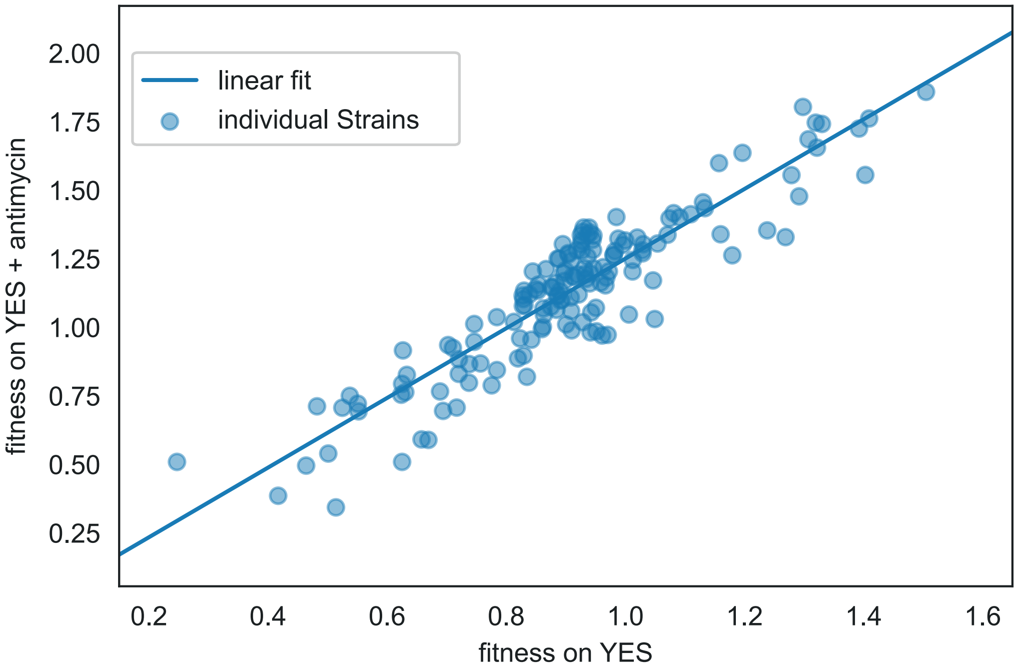


Supplementary Figure 1: Fitness (corrected colony size relative to standard lab strain 972) of wild strains on YES and YES with 500ug/L antimycin A. A linear regression with y = 1.27x - 0.02 describes this relationship well ($R^2 = 0.83$) which means most of the variation of fitness on YES + antimycin is explained by basal growth rate and not specific to the effect of antimycin. Antimycin resistance is determined by taking the ratio of the fitness of YES + antimycin and YES alone.

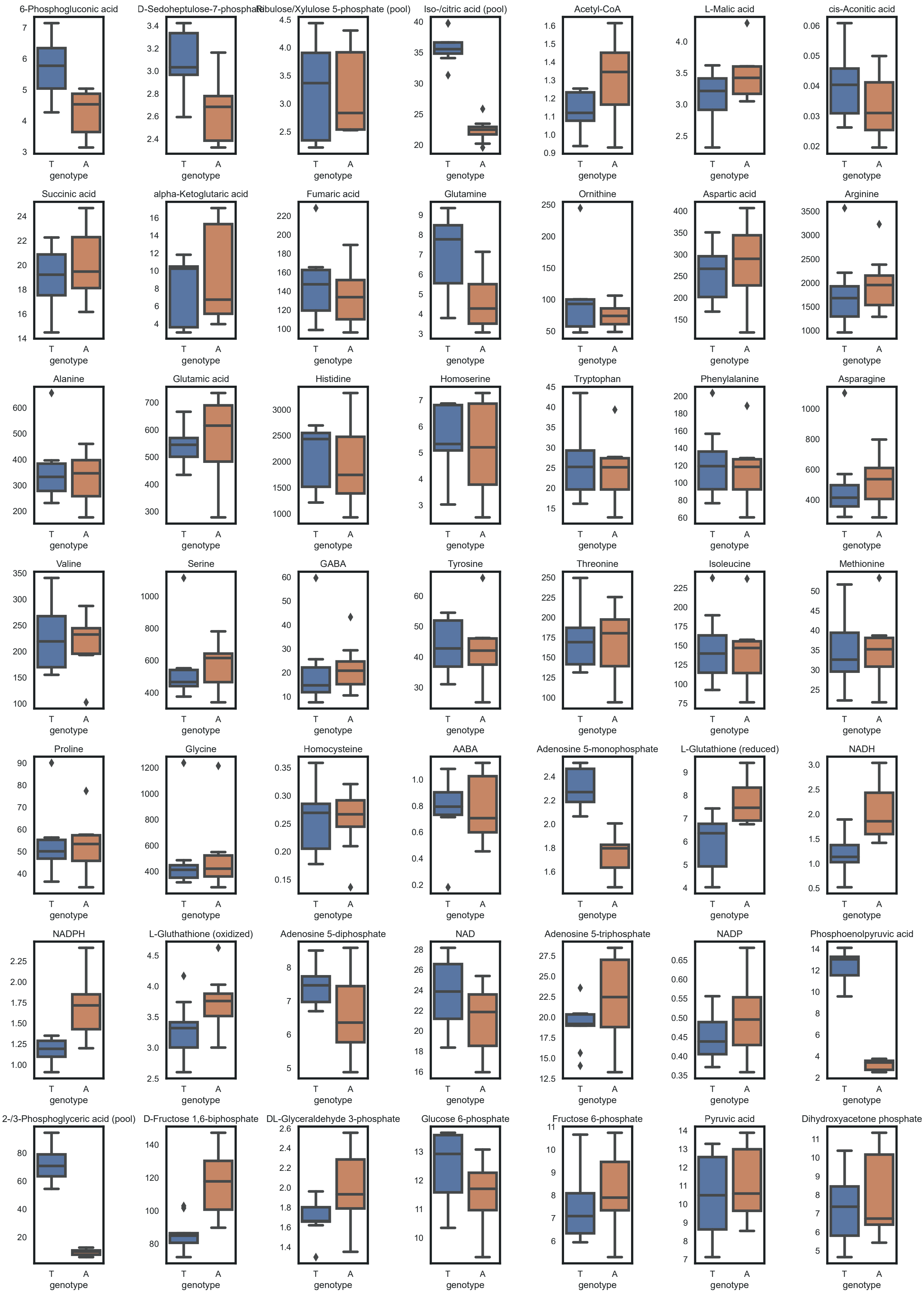


Supplementary Figure 2: Boxplots comparing intracellular metabolite concentrations (concentrations in the prepared sample divided by the optical density of the culture at sampling) of the T-strain (*S. pombe* reference strain *968*) and A-strain (*968 pyk1*$^{T343A}$).

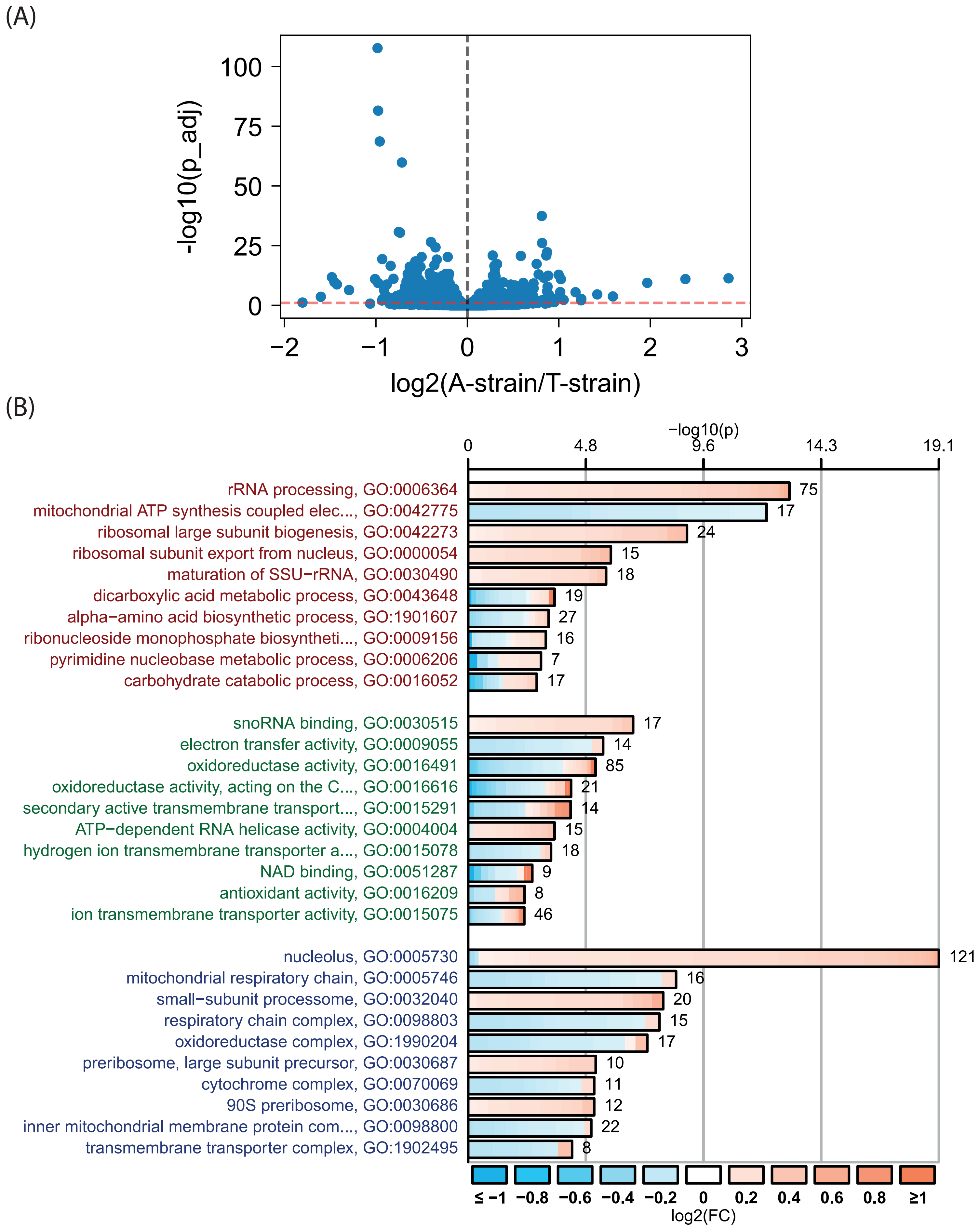


Supplementary Figure 3: (A) Volcano plot of transcriptome data set. (B) CellPlots summarising GO enrichment analysis of transcripts differentially expressed in A-strain versus T-strain.

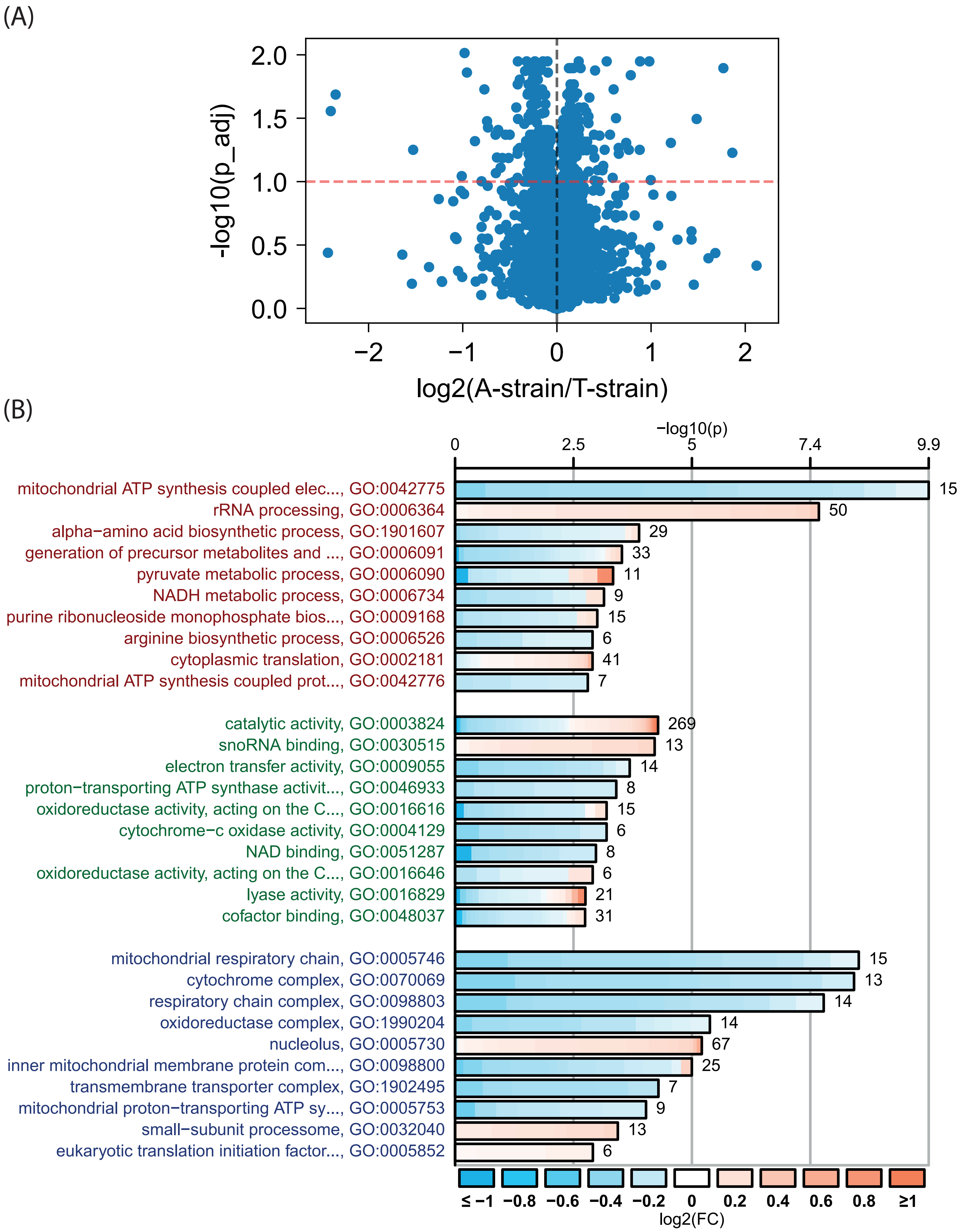


Supplementary Figure 4: (A) Volcano plot of transcriptome data set. (B) CellPlots summarising GO enrichment analysis of proteins differentially expressed in A-strain versus T-strain.

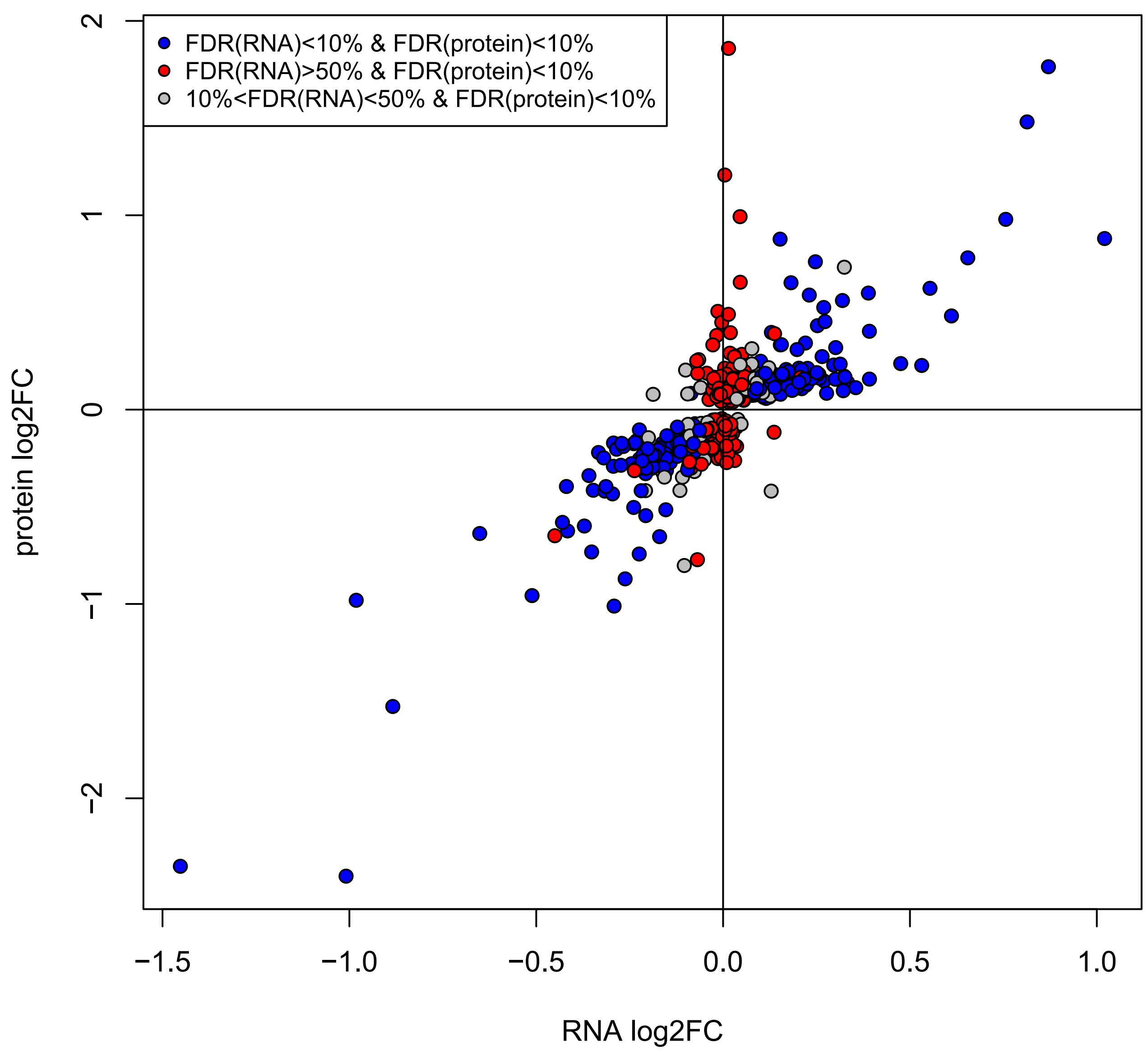


Supplementary Figure 5: Differential abundance of transcripts and proteins. All genes with differential protein abundance between the A-strain and the T-strain at FDR<10% are shown. Genes for which the transcripts were not deferentially abundant (FDR>50%) are shown in red. Genes whose transcripts and proteins were deferentially abundant at FDR<10% are shown in blue. Fold-changes were computed as log2(A)-log2(T). Genes that were only affected on the protein level (red) were significantly enriched in functions related to cytoplasmic translation and depleted in genes involved in ribosome biogenesis compared to genes affected on both levels (blue, Supplementary Tables 5 and 6).

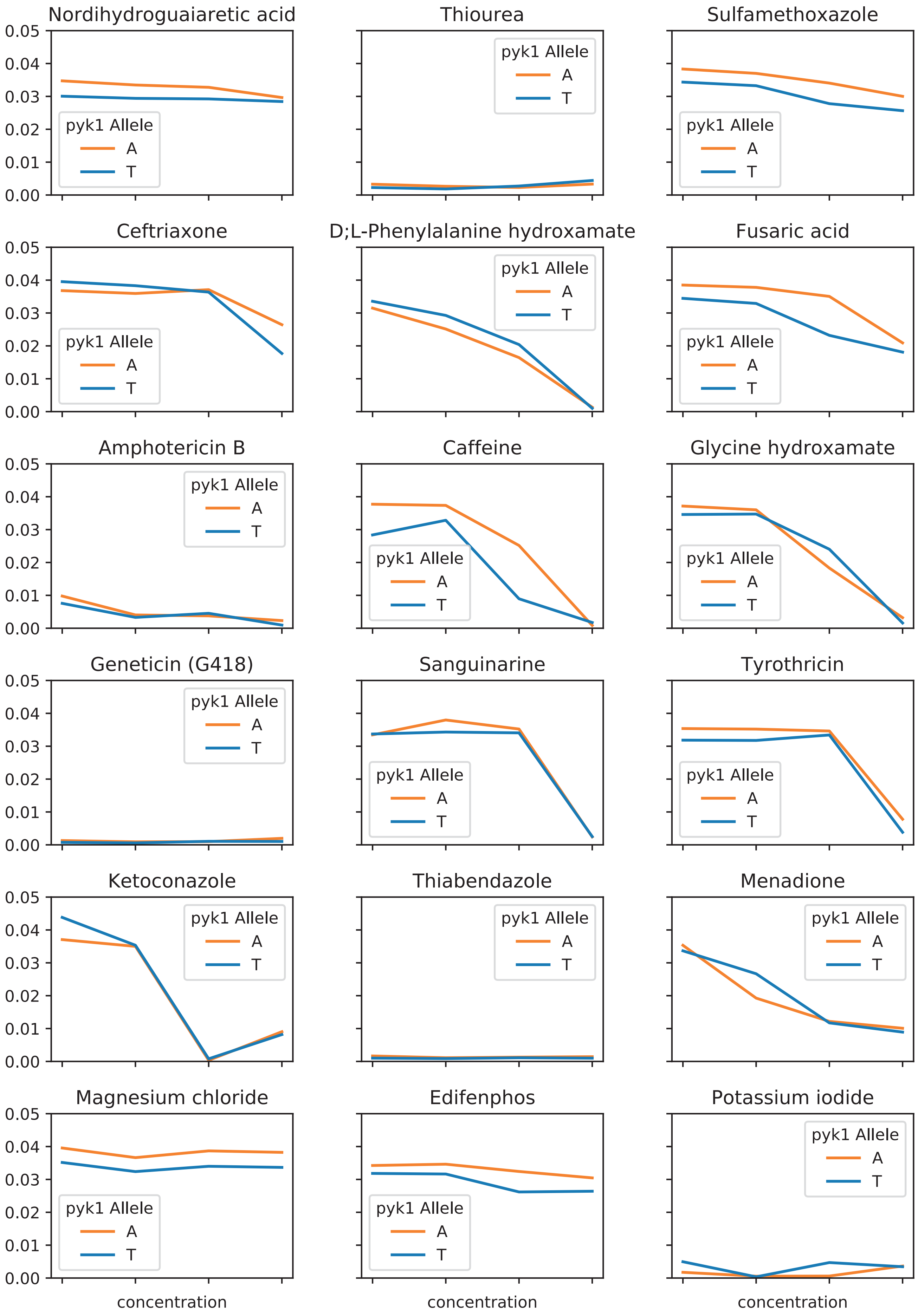


Supplementary Figure 6 (Page 1): Dose-response curves of T- and A-strain, showing maximum growth rate (y-axis) for each of the 4 concentration levels contained in the Biolog Phenotype Array.

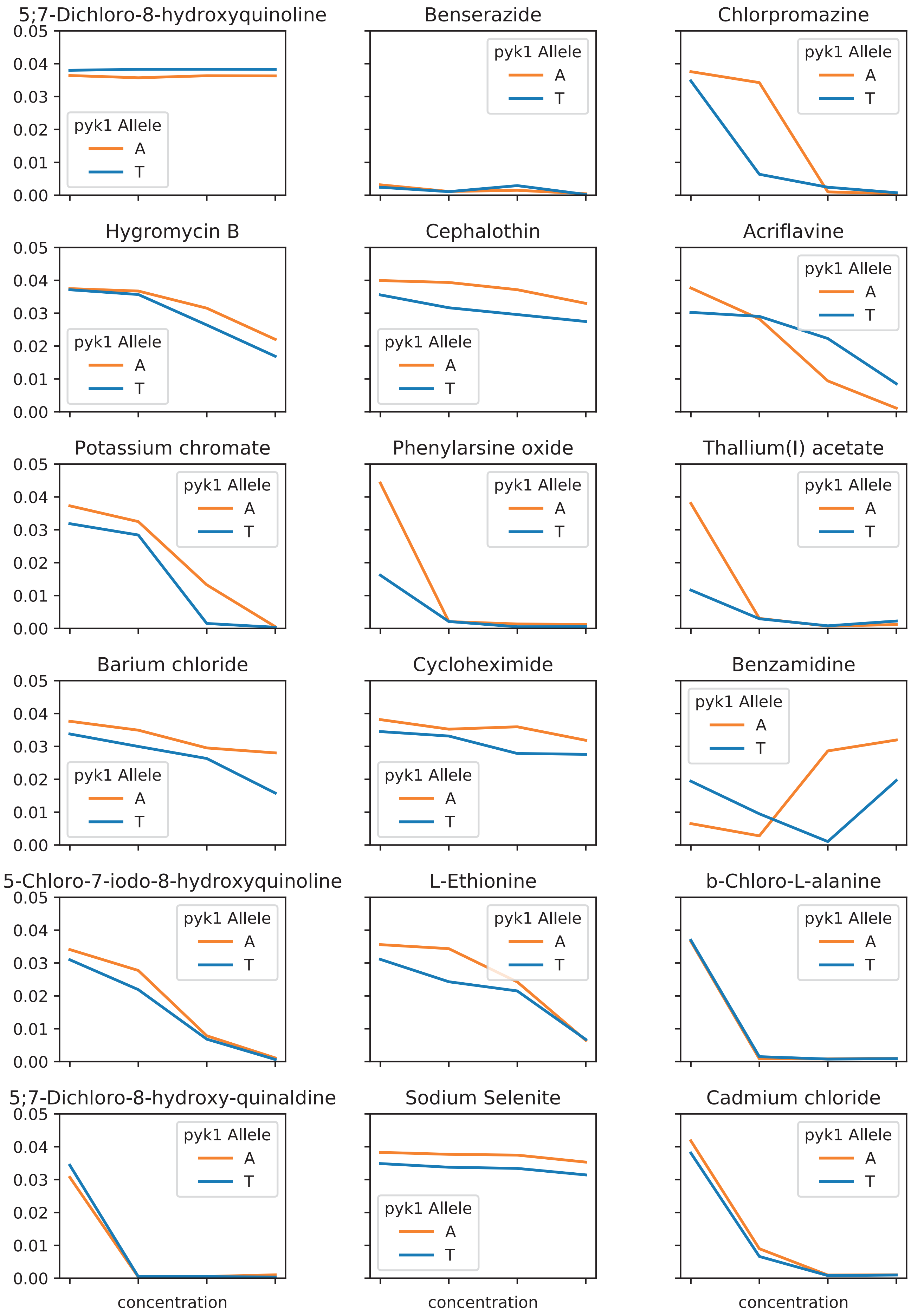


Supplementary Figure 6 (Page 2): Dose-response curves of T- and A-strain, showing maximum growth rate (y-axis) for each of the 4 concentration levels contained in the Biolog Phenotype Array.

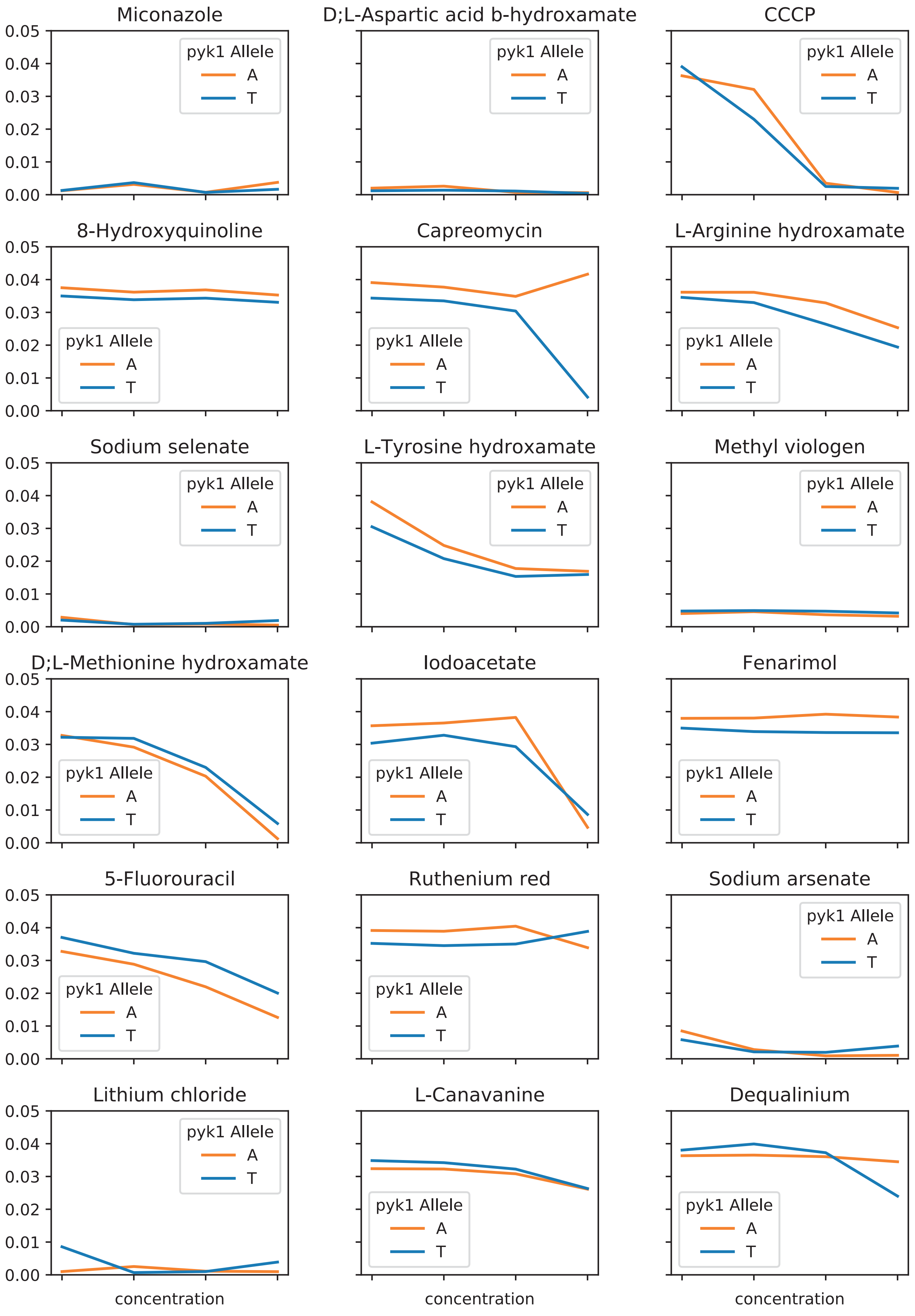


Supplementary Figure 6 (Page 3): Dose-response curves of T- and A-strain, showing maximum growth rate (y-axis) for each of the 4 concentration levels contained in the Biolog Phenotype Array.

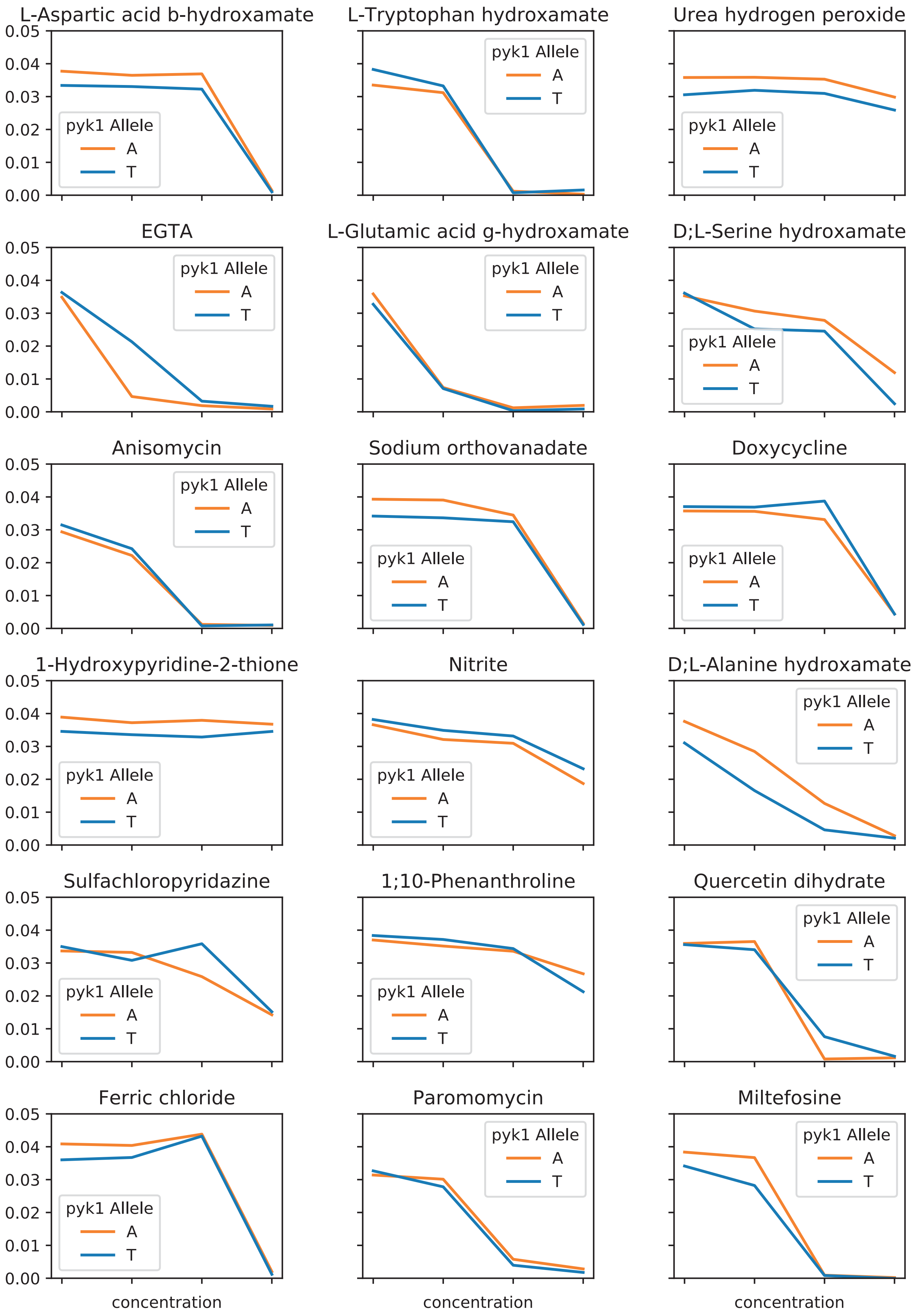


Supplementary Figure 6 (Page 4): Dose-response curves of T- and A-strain, showing maximum growth rate (y-axis) for each of the 4 concentration levels contained in the Biolog Phenotype Array.
